# Functions of olfactory receptors are decoded from their sequence

**DOI:** 10.1101/2020.01.06.895540

**Authors:** Xiaojing Cong, Wenwen Ren, Jody Pacalon, Claire A. de March, Lun Xu, Hiroaki Matsunami, Yiqun Yu, Jérôme Golebiowski

## Abstract

G protein-coupled receptors (GPCRs) conserve common structural folds and activation mechanisms, yet their ligand spectra and functions are highly diversified. This work investigated how the functional variations in olfactory GPCRs (ORs)-the largest GPCR family-are encoded in the primary sequence. With the aid of site-directed mutagenesis and molecular simulations, we built machine learning models to predict OR-ligand pairs as well as basal activity of ORs. *In vitro* functional assay confirmed 20 new OR-odorant pairs, including 9 orphan ORs. Residues around the odorant-binding pocket dictate the odorant selectivity/specificity of the ORs. Residues that encode the varied basal activities of the ORs were found to mostly surround the conserved motifs as well as the binding pocket. The machine learning approach, which is readily applicable to mammalian OR families, will accelerate OR-odorant mapping and the decoding of combinatorial OR codes for odors.

## Introduction

Functions of proteins are encoded within either diversified or conserved subparts of their sequence. G protein-coupled receptors (GPCRs) are the most remarkable examples of this phenomenon. GPCRs are the largest membrane protein family and the targets of about 40% of marketed drugs^1^. The human genome contains over 800 genes coding for GPCRs, which exert differentiated and specific functions in the complex cellular signaling network. Half of these genes are olfactory receptors (ORs), which endow us with fascinating capacities of odor discrimination^2,3^. Mammalian GPCRs conserve a typical structural architecture of seven transmembrane helices (7TM) that house an orthosteric ligand-binding pocket^4^. Their intrinsic signaling mechanism, large-scale conformational changes to accommodate their cognate G proteins, is encoded in conserved motifs throughout the 7TM, which form a network of inter-TM contacts converging to their cytoplasmic part^5^. Namely, the “D(E)RY”, “CWLP” and “NPxxY” motifs in TM3, TM6 and TM7, respectively, are the most conserved hubs of the allosteric communication between the orthosteric pocket and the cytoplasmic side of class A GPCRs^4^. The rest of the GPCR sequences, especially the ligand-binding pocket, have diversified extensively, which resulted in huge variations in the receptors’ function. Amongst the hundreds of variable residues in the GPCR sequences, it is very likely that some are specialized for the receptors’ ligand specificity/selectivity while, others encode more information on the receptors’ intrinsic activity.

This study focused on the divergence of ligand-dependent and ligand-independent (or basal) activities of olfactory class A GPCRs. We seek to identify amino acid positions that are critical for the receptors’ functional heterogeneity. ORs discriminate a vast spectrum of volatile molecules (odorants) and code for an innumerous number of odors perceived in the brain. The many-to-many relationships between ORs and odorants are key to understanding odor perception^6^. ORs are also expressed ectopically, some of which have emerged as potential drug targets^7–9^. To date, ~45% of the human ORs (hORs) and ~10% of the mouse ORs (mORs) have been deorphanized with sometimes only one but also with more odorants (Table S1)^10–16^. We naturally hypothesize that ligand selectivity and specificity are largely dictated by the binding affinity which is encoded in the binding pocket^17,18^. Indeed, ORs responding to the same odorants share higher sequence homology around the pocket than in the rest of the receptor^10^. Yet, affinity does not correlate with efficacy nor potency. GPCR response to ligands is a complex phenomenon that can be drastically altered by mutations far-off the pocket^19^. Basal activity may play an important role therein, including for ligand selectivity^20^. We have previously found that OR basal activity is partly correlated with the broadness of the ORs’ ligand spectrum^21^. Differential OR basal activities have been suggested to mediate OR neuron axon targeting to specific glomeruli in the olfactory bulb, although clear evidence is still lacking^20^. The basal activity is associated with the intrinsic activation mechanism and is likely encoded around the highly conserved hubs controlling this mechanism.

Machine learning has shown success in protein function predictions using amino-acid sequences (reviewed in ref.^22^). Sequence-based approaches circumvent the difficulties to obtain high-resolution structures for some proteins, which is the case of ORs. However, detailed functional descriptions such as ligand selectivity/specificity are challenging, due to the complex nature of the problems and the lack of data. Encouraging studies have been made on ORs owing to their large number of known ligands^10,23^. However, in these systems the signal-to-noise ratio is low. If one assumes that a given function is mostly encoded into 20 residues, there are more than 10^30^ possibilities of selecting 20 residues out of a GPCR sequence of ~300 residues. Therefore, the selection of relevant residues is key to constructing accurate machine learning models. A recent study on insect and mammalian ORs consistently demonstrated that selection of subsets of 20 residues could significantly increase the signal-to-noise ratio^24^. Here, we used molecular modeling and site-directed mutagenesis data to guide this selection. Random forest (RF) and Recursive partitioning and regression trees (RPART) were then employed to predict the agonist-stimulated and the basal activity of human, mouse and some primate ORs, based on specific residues. *In vitro* functional assays assessed the predictivity of the machine learning models. This approach (outlined in Fig. 1A and detailed in Materials and Methods) may be applicable to other protein families and functions.

**Fig. 1.**
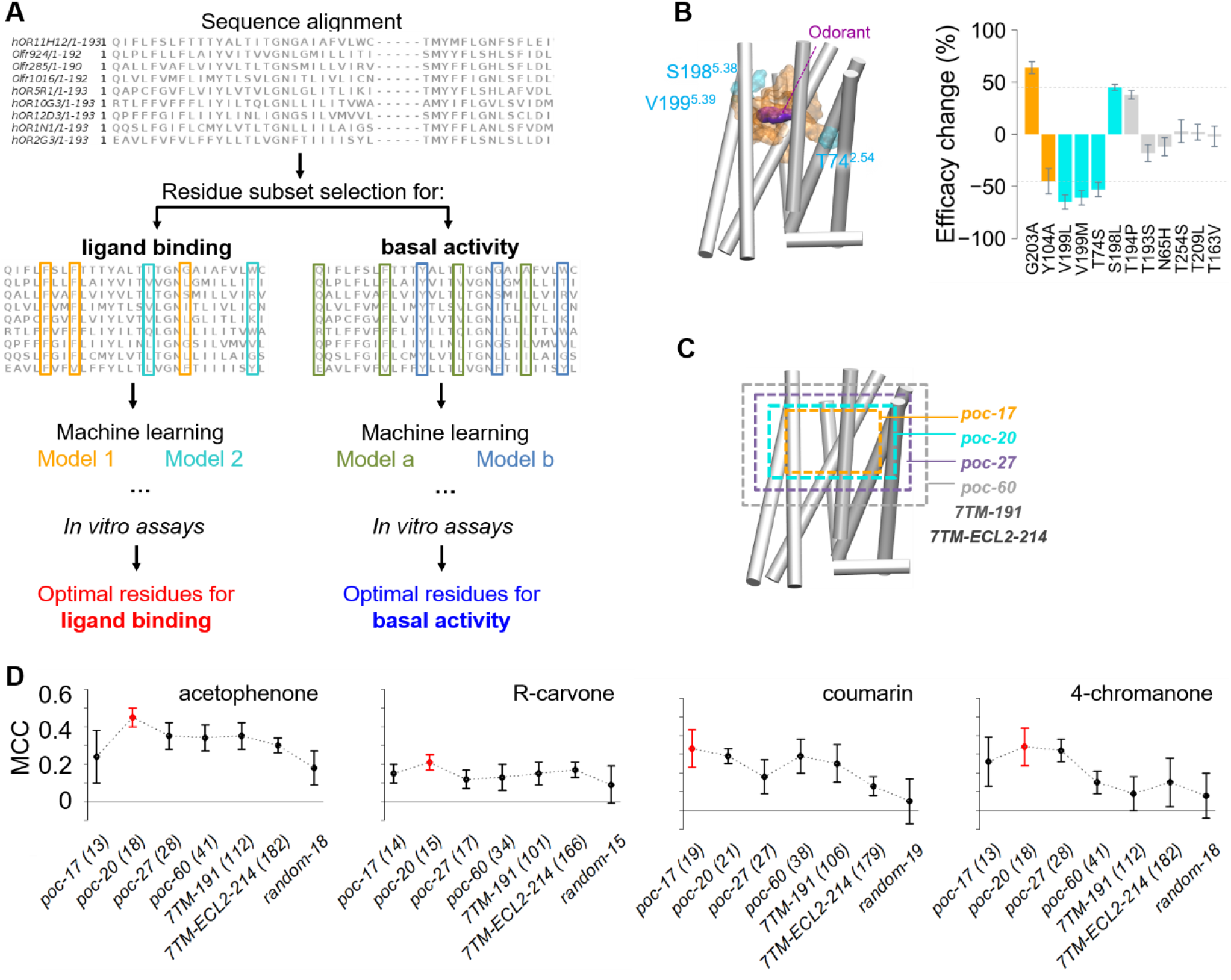
Machine-learning protocol and selection of key residues by molecular modeling and site-directed mutagenesis.

**(A)** Machine learning workflow, where different residue subsets were extracted from the sequence alignment for the training of different models for either ligand binding or basal activity prediction. The performance of each model was assessed *in vitro*, to identify the optimal residue subsets that encode each function. **(B)** 3D model of mOR256-31 illustrating the 17 odorantcontacting residues (orange) and V199^5.39^, T74^2.54^ and S198^5.38^ (cyan) that were newly identified as part of the binding pocket. Bar plot shows the effect of point mutations on the odorants’ efficacy on mOR256-31 (see Fig. S1 for the dose-dependent response curves). Mutations at G203^5.43^ and Y104^3.32^ (orange bars) were used as positive controls since they are known to be crucial for binding coumarin and R-carvone. Cyan bars represent residues that significantly affected the efficacy of R-carvone on mOR256-31. **(C)** Outline of the 6 subsets of residues used for feature selection: i) *poc-17*, the 17 odorant-contacting residues; ii) *poc-20*, the 20 pocket residues; iii) *poc-27*, an extension of the pocket region (27 residues) up to a 6 Å distance around the odorants; iv) *poc-60*, a larger extension (60 residues) until 8 Å from the odorants; v) *7TM*, the 7TM bundle consisting of 191 residues; vi) *7TM-ECL2-214, 7TM-191* plus residues in ECL2 between the two conserved cysteines, increasing the number of residues to 214. Other loop regions were excluded because they contain many gaps in the sequence alignment. **(D)** MCC of the RF classifiers during 5-fold cross-validation repeated 5 times. Labeled in parenthesis is the number of optimal residues that gave the best performance in each model. Error bars indicate SEM.

## Results

### Feature selection for machine learning of OR response to odorants

Four odorants, acetophenone, coumarin, R-carvone, and 4-chromanone, were considered. They are chemically related and may not represent the odorant space. We chose these odorants because dozens to hundreds of OR responses were available to feed the machine learning algorithms (Table 1 and Table S2). Using the sequence alignment of their cognate ORs, we trained classification models that read the sequence of new ORs and predicted whether they responded to the four odorants.

**Table 1.**
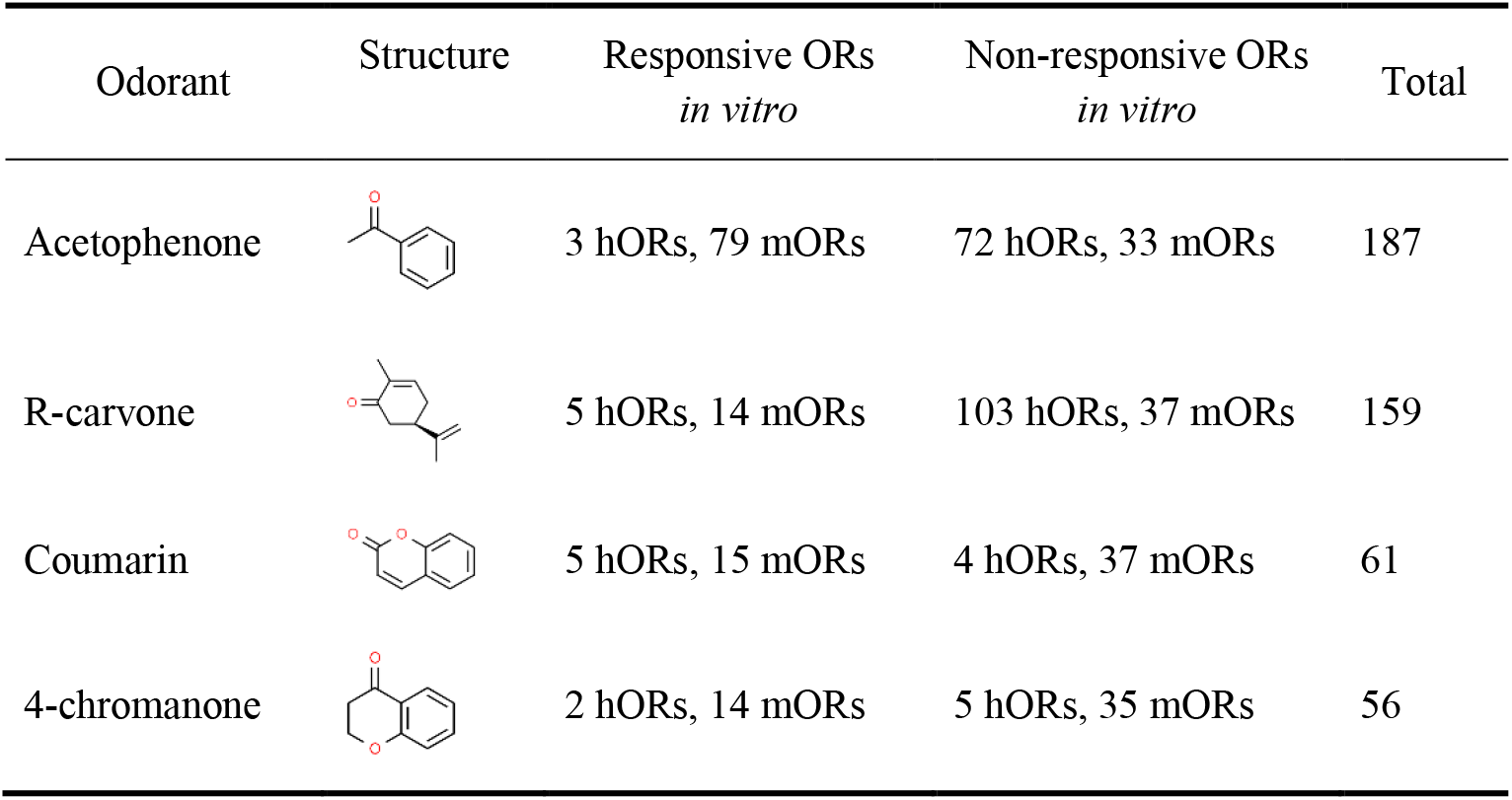
Literature data of human and mouse ORs tested against the 4 odorants. See Table S2 for the full list.

| Odorant | Structure | Responsive ORs<br>*in vitro* | Non-responsive ORs<br>*in vitro* | Total |
| --- | --- | --- | --- | --- |
| Acetophenone |  | 3 hORs, 79 mORs | 72 hORs, 33 mORs | 187 |
| R-carvone |  | 5 hORs, 14 mORs | 103 hORs, 37 mORs | 159 |
| Coumarin |  | 5 hORs, 15 mORs | 4 hORs, 37 mORs | 61 |
| 4-chromanone |  | 2 hORs, 14 mORs | 5 hORs, 35 mORs | 56 |

<u>Rational selection of most relevant residues</u>. Intuitively, we focused on the odorant-binding residues within the canonical ligand-binding pocket. To identify them, molecular modeling and site-directed mutagenesis were performed on mOR256-31 (Olfr263), a broadly tuned receptor considered as prototypical because it responds to three of the four odorants (coumarin, R-carvone and acetophenone)^21,25^. Previously established approaches^19,21,26^ were used to construct 3D models of mOR256-31 bound with the odorants. The model was built under the constraint of site-directed mutagenesis data covering 96 positions throughout the TM bundle. Molecular dynamics (MD) simulations illustrated 17 residues within a 5 Å distance of the bound odorants (Table S3), which were considered as a starting point for machine learning. Fourteen of these residues had been shown by site-directed mutagenesis to be important for the response of one or multiple ORs to their ligands (Table S3). We additionally mutated residues around the odorants positions and identified three additional residues where mutations significantly affected the ligand efficacy *in vitro* (cyan bars in Fig. 1B). Therefore, 20 residues were considered as a more accurate binding pocket (named *poc-20* hereafter), including 17 in direct contact with the odorants (named *poc-17*).

To evaluate whether the above-identified pocket residues were the most relevant to predict ligand binding, we tested 6 small-to-large residue subsets as heuristics (*poc-17, poc-20*, as well as *poc-27* and *poc-60, 7TM-191, 7TM-ECL2-214*, see Fig. 1C and Table S4). These were considered as initial guesses of relevant residues for model training (Fig. 1A) on 80% of the training set data. The remaining 20% were used for model validation (the validation set). Starting from these initial guesses, two steps of feature selection (see Materials and Methods) were carried out until the model converged to an optimized set of residues. This procedure allowed searches for optimal residues around the initial guess to be performed and was crucial in this evaluation as it is unrealistic to test all possible residue subsets through a brute force approach. Using the optimized residue subsets, we trained RPART, SVM and RF classifiers and compared their predictivity on the validation set of data, which was measured by Matthew’s correlation coefficient (MCC)^27^. RF performed the best amongst the three algorithms. The performance analysis on the validation set revealed that the best RF classifiers for acetophenone, R-carvone and 4-chromanone were initialized with *poc-20*, whereas the one for coumarin was initialized with *poc-17* (Fig. 1D). They converged on 15-21 residues after optimization, depending on the odorant (Fig. 1D). Control models built with the same number of randomly selected residues showed nearly no predictivity (Fig. 1D). We then used the same procedure with 100% of the data for each odorant to construct four optimal RF classifiers. These classifiers screened ~600 OR sequences and predicted their probability to respond to each of the 4 odorants.

#### In vitro assessment of machine learning predictions

We predicted the probability of 605 ORs to respond to the 4 odorants. The ORs comprise the entire hOR family including some genetic variants (498 sequences), as well as 107 mORs for which the plasmids were available in our laboratory. They were ranked by the mean value of their 4 probabilities. We chose 37 top-ranked ORs for *in vitro* functional assays, which were predicted to respond to at least one odorant. As negative controls, we selected 6 ORs among those predicted to be non-responsive to any of the odorants. The total of 43 ORs were tested against all the 4 odorants (Fig. 2A). When significant responses were observed at 300 μM, dose-dependent responses were measured.

**Fig. 2.**
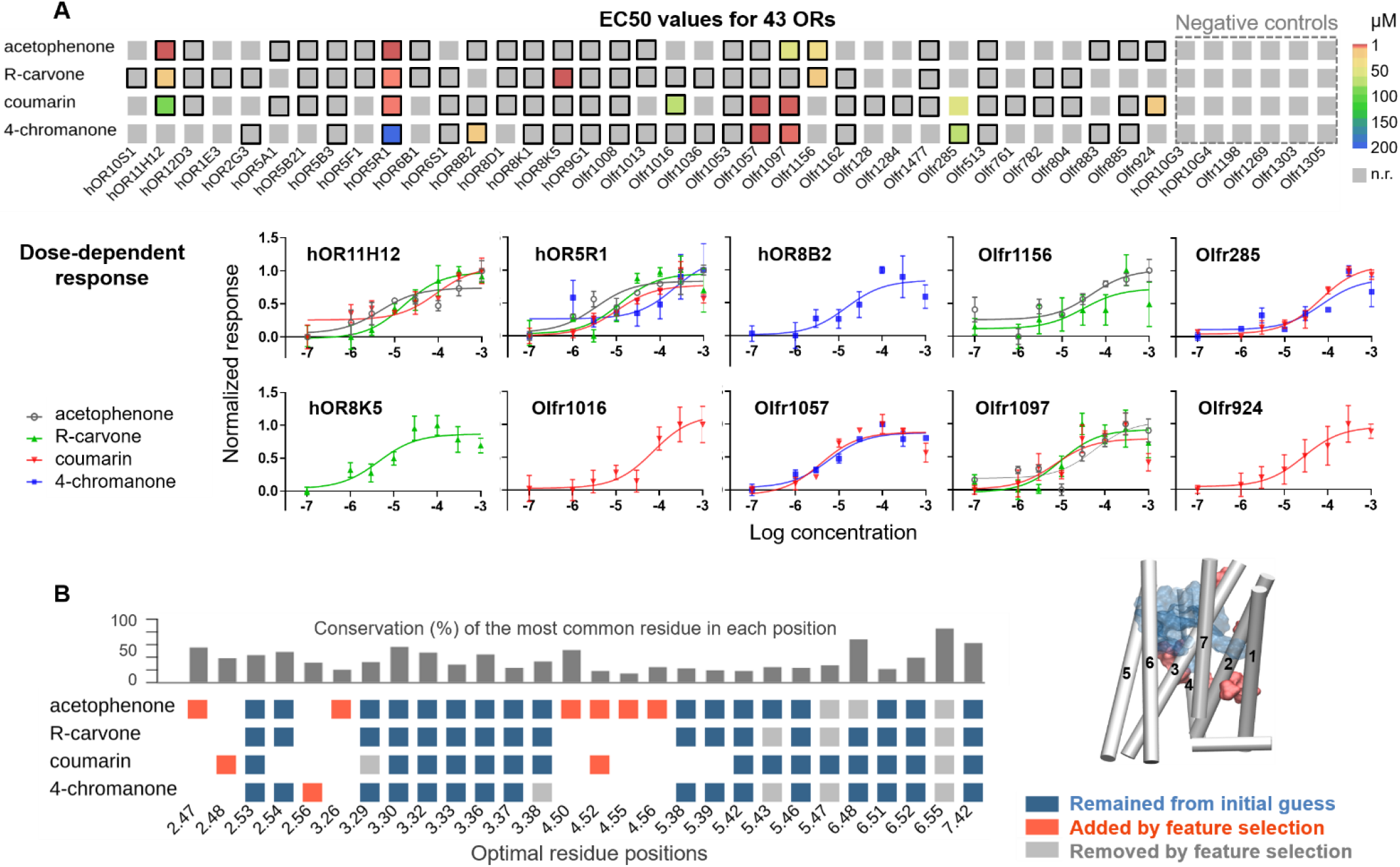
Machine learning model performance evaluated with external test sets from *in vitro* functional assays.

**(A)** Heatmap of EC50 values and dose-dependent response curves. Predicted OR-odorant pairs are boxed in black. The 6 negative-control ORs are indicated with gray dashed line. **(B)** Ballesteros-Weinstein numbering of the residues that produced the most predictive model for each odorant. Dark blue, red and light gray indicate the residues that were retained, added or removed by the feature selection procedure, respectively. Bar plot on top shows the conservation of the most common residue in the combined hORs and mORs alignment.

Ten ORs responded to at least one of the odorants in a dose-dependent manner, resulting in 20 new OR-odorant pairs with EC50 values ranging from 3 μM to 218 μM (Fig. 2A, Table 2 and Table S5). Nine ORs (3 hORs and 6 mORs) were deorphanized here, including 6 that responded to multiple odorants. The 6 negative-control ORs showed no significant response to any of the odorants. We checked that the non-responsive ORs were expressed in the cell membranes with measurable basal activities. Seven of them including 2 negative controls (hOR1E3, hOR5A1, hOR5B1, hOR8D1, hOR9G1, hOR10G3 and hOR10G4) had been previously shown to respond to other odorants in the same heterologous cell functional assays^12^. Note that the lack of response of the other non-responsive ORs might be due to impaired functions in the heterologous cells. This remains to be assessed when these ORs will be deorphanized. The RF classifier showed 14%–25% hit rates for the 4 odorants (Table 3). The predictivity (MCC) ranged from 0.18 to 0.32 (Table 3). Control models built with randomly selected residues consistently showed near zero predictivity (Table 3). We also tested other residue subsets but none of them outperformed the RF classifier used in the virtual screening (Fig. S2). *In vitro* data confirmed that for all the 4 odorants, the residues in the optimal RF classifiers contained the highest signal-to-noise ratio. The optimized residue subsets are intuitively located around the binding pocket, most of which belong to the initial guesses, *poc-17* (for coumarin) and *poc-20* (Fig. 2B). These residues show very low conservation in hORs and mORs (Fig. 2B), suggesting that they have diversified to differentiate diverse ligands^17,18^

**Table 2.** New OR-odorant pairs verified by *in vitro* dose-dependent response^*a*^

| Odorant | OR gene |
| --- | --- |
| R-carvone | hOR11H12, hOR5R1, hOR8K5, Olfr1156 |
| Coumarin | hOR11H12, hOR5R1, Olfr924, Olfr285, Olfr1016, Olfr1057, Olfr1097 |
| 4-chromanone | hOR5R1, hOR8B2, Olfr1057, Olfr1097, Olfr285 |
| Acetophenone | hOR11H12, hOR5R1, Olfr1097, Olfr1156 |

<sup>a</sup> One-way Anova with Benjamini - Hochberg FDR correction.

**Table 3.**
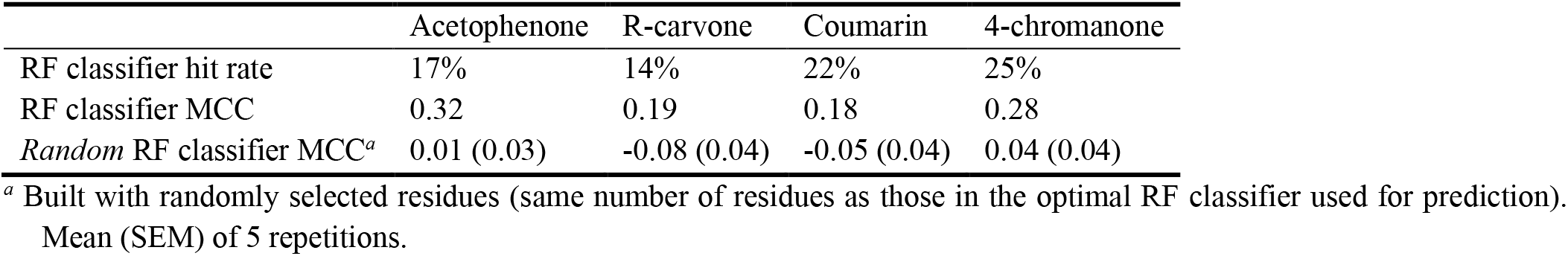
Performance of the RF classifier in predicting new ORs.

|  | Acetophenone | R-carvone | Coumarin | 4-chromanone |
| --- | --- | --- | --- | --- |
| RF classifier hit rate | 17% | 14% | 22% | 25% |
| RF classifier MCC | 0.32 | 0.19 | 0.18 | 0.28 |
| Random RF classifier MCC<sup>a</sup> | 0.01 (0.03) | -0.08 (0.04) | -0.05 (0.04) | 0.04 (0.04) |

<sup>a</sup> Built with randomly selected residues (same number of residues as those in the optimal RF classifier used for prediction). Mean (SEM) of 5 repetitions.

### Prediction of OR basal activity

The basal activity relates to the intrinsic activation mechanism, itself encoded in the highly conserved residues in GPCRs. It had been frequently reported that mutations within and around GPCR conserved motifs affect receptor basal activity remarkably (Table S6, Fig. S3). Therefore, the selection of residue subsets for machine learning of OR basal activity was centered on the conserved motifs. We selected residues in a 4 Å distance around the conserved motifs of ORs, which were identified by structural analyses of 3D models of mOR256-3, mOR256-8 and mOR256-31. This subset was made up of 49 residues and named *res-49*. To produce alternative and larger subsets, site-directed mutagenesis was performed on mOR256-8 to identify residues that affect its basal activity. We mutated 72 residues around the conserved motifs and measured if they have a major, moderate, or minor impact on the receptor’s basal activity (Fig. 3A).

**Fig. 3.**
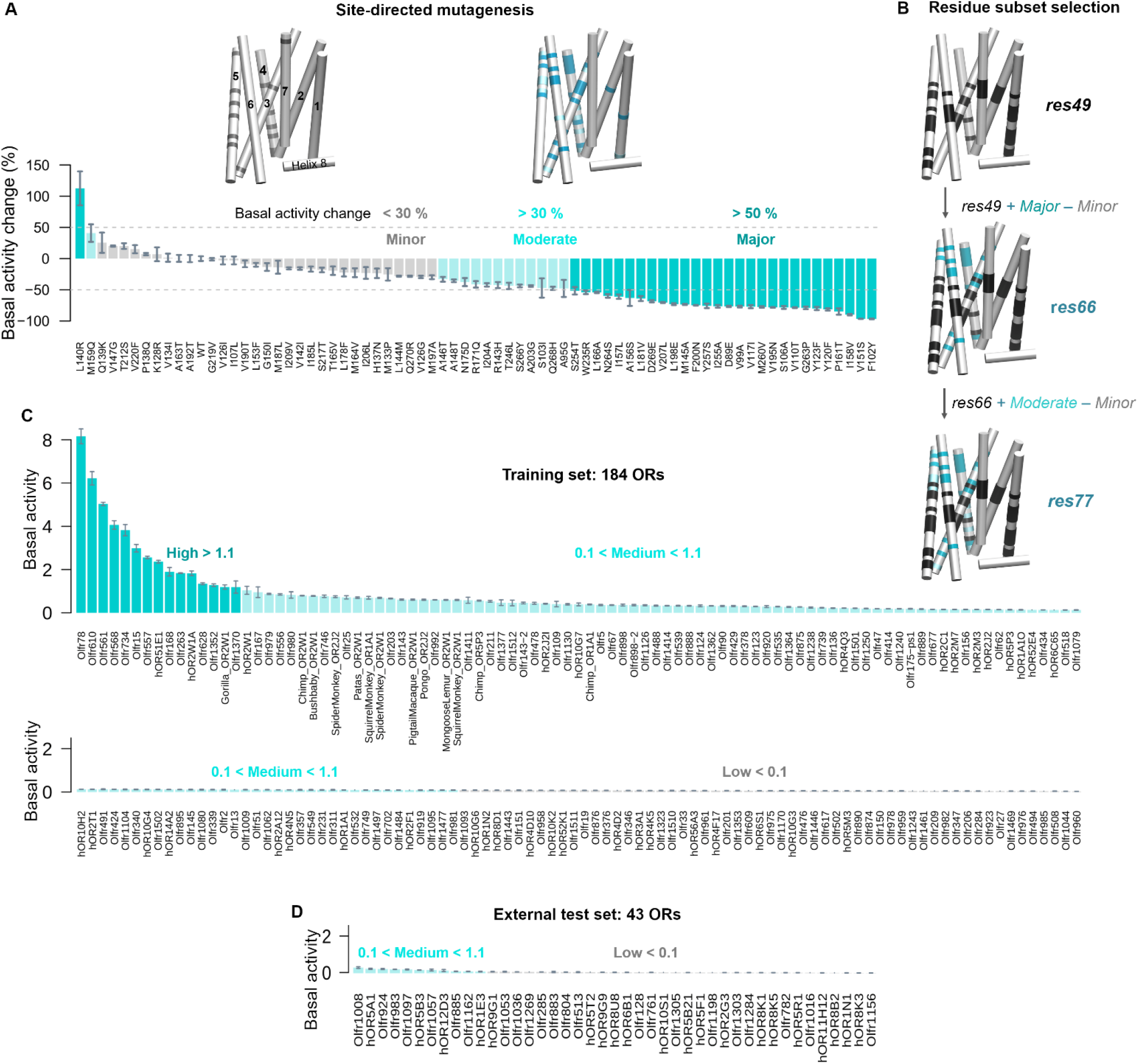
*In vitro* basal activities.

**(A)** Effects of 72-point mutations on the basal activity of mOR256-8 *in vitro*. Plotted are percentage changes with respect to the wild type (WT). We verified that the changes were not due to altered expression levels. **(B)** Selection of residue subsets based on *res-49* (black, residues in 4 Å distance around the conserved motifs of ORs). **(C)** *In vitro* basal activity of the training set of 184 ORs for machine learning, divided in 3 classes. **(D)** Basal activity of 43 ORs to assess the prediction of the optimal RPART classifier. Error bars indicate SEM (*n* = 3–9, except for hOR2W1 where n = 39).

Like the machine learning procedure above, we tested five heuristic subsets (*res-49, res-66, res-77, 7TM-191* and *7TM-ECL2-214*) as initial guesses for model training. *res-66* and *res-77* were extensions of *res-49* by adding residues with “major” and “moderate” impacts on mOR256-8 basal activity, accumulatively (Fig. 3B). The same modeling-training procedure as above (Fig. 1A) was applied to construct RF, RPART and SVM classifiers. The data for model training was generated by *in vitro* measurements of the basal activity of 183 mammalian ORs, 108 of which were recently published^11^ (Fig. 3C). The models were trained to classify the basal activities of candidate ORs into 3 classes (low, medium and high, Fig. 3A). Cohen’s kappa statistic^28^ handles multi-class problems and was used as benchmark for the predictivity of each model trained on 80% of the data and tested on the remaining 20%. The RPART classifier initialized with *res-66* was optimized to 32 residues and gave the best results on the validation set (Fig. S4A). This choice was then used to build an optimal RPART classifier with 100% of the data.

Using the optimal RPART classifier, we predicted and measured the basal activities of 43 new ORs (Fig. 3D). The model showed Cohen’s kappa of 0.29, in agreement with the predictivity on the training set during 5-fold cross validation (Fig. S4A). Control models built with randomly selected residues or the other residue subsets showed lower predictivity (Fig. S4B), as expected. Thus, the optimal classifier was initialized with *res-66*, which finally converged to 30 residues (Fig. 4A). These residues surround the conserved motifs and the ligand-binding pocket, including one in the ECL2 next to a highly conserved disulfide bond (Fig. 4A). A recent study indeed reported roles of ECL2 in modulating class A GPCR basal activity^29^.

**Fig. 4.**
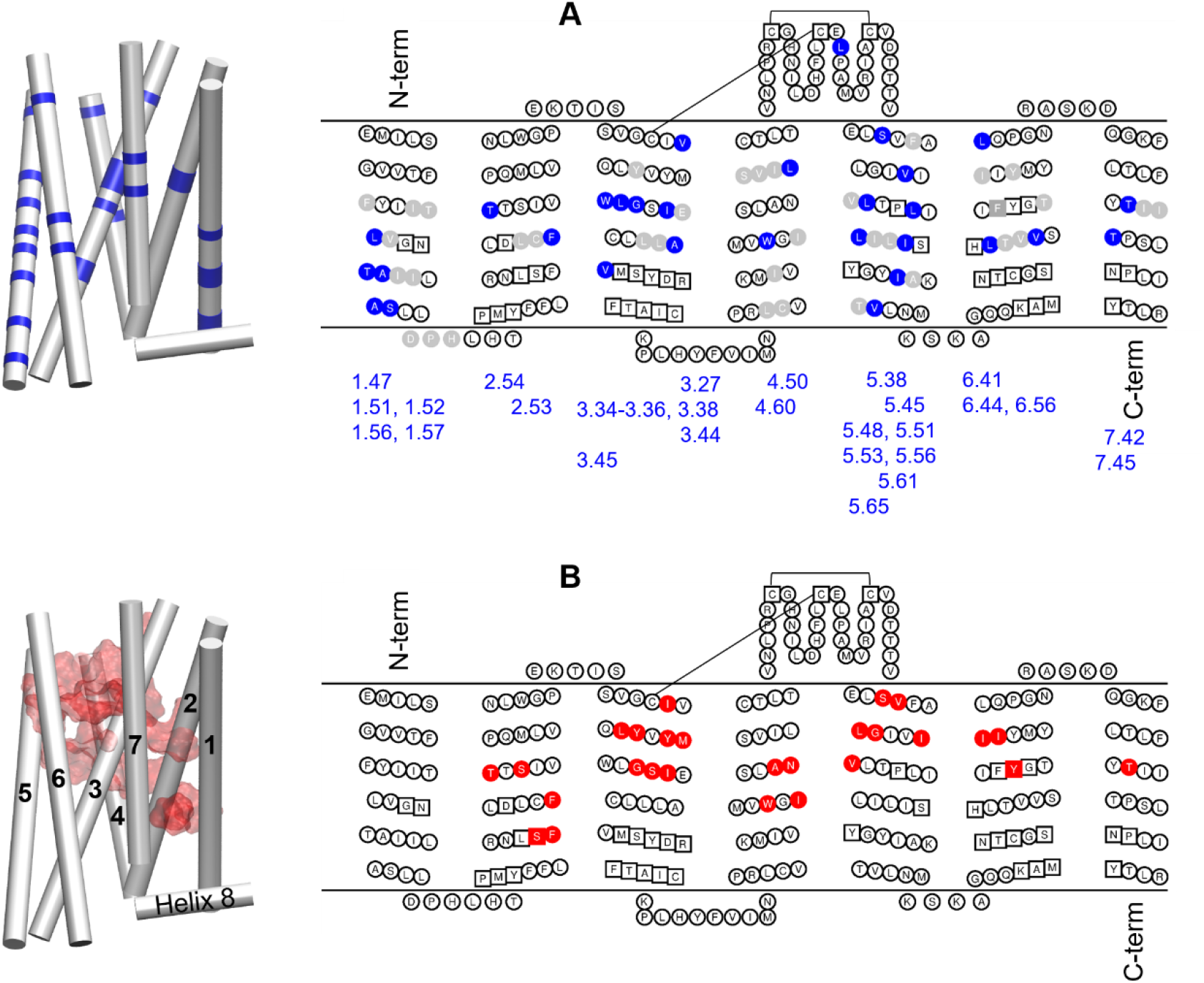
Location of optimal residue subsets illustrated with mOR256-31.

3D homology model of 7TM and snake plot illustrating the locations of the optimal residue subsets for basal activity (blue) **(A)** and ligand binding (red) **(B)**, respectively. Conserved motifs in ORs are squared. Light gray indicates the residues removed from the initial guess (*res-66*) by feature selection. The N-and C-termini are truncated for clarity.

## Discussion

This study illustrated how the ORs’ functional heterogeneity is encoded in their sequence, through *in* vitro assays, structural modeling, and machine learning. We measured the basal activity of 299 mammalian ORs (108 mORs published in our previous work and 72 mutants of mOR256-8) as well as the activity of 4 odorants on 59 ORs (incl. 16 mutants of mOR256-31). The machine learning feature selection allowed for searches of most relevant residues, initialized with knowledge from structural modeling and site-directed mutagenesis. We identified 17-20 residues around the odorant-binding pocket which differentially dictate the variation in the ORs’ response to the odorants (Fig. 4B where all residues cumulate to 27). The ligand-binding pocket of GPCRs have evolved with high diversity to discriminate various stimuli. It is not surprising that the ORs’ response to the odorants could be predicted to a great extent with less than 10% of the sequence, made up with highly variable residues. The results validate previous predictions of pocket residues based on OR sequence analysis^17,18^ and numerous site-directed mutagenesis data^18,19^, which are located in the upper portion of TM3, 5, 6, and 7. Here we highlighted 5 additional residues in TM2, surrounding a conserved allosteric site (centered at D^2.50^). The allosteric site in non-olfactory class A GPCRs (typically composed of D^2.50^, N^3.35^ and S^3.39^) is known to bind the Na^+^ ion, which modulates the receptor’s activation and affinity/response to ligands (reviewed in ref.^30^). Most ORs contain a second acidic residue (Ballesteros-Weinstein number 3.39) at this site, which might also accommodate divalent cations^30^. While copper ions play important roles in the recognition of sulfur odorants^31,32^, it remains unclear whether this conserved site in the ORs is involved. The machine learning protocol established here represents an alternative to existing models using full sequences^10,23^. It teased out residues that are specific for each odorant, which is essential for understanding how chemically similar odorants are differentiated by the OR family with high specificity/selectivity. So far, research focusing on subtle selectivity has mostly employed site-directed mutagenesis and molecular modeling of individual ORs, namely hOR17-4 for lilial and bourgenoal analogues^33^, hOR1A1 for R-/S-carvone enantiomers^34^, hOR5AN1 and mOR215-1 for musk odorants^35^, mOR-EG for eugenol^36^, and our recent study on zebrafish ORs for bile acids/salts^37^.

Regarding the basal activity, we found that the variation is mostly governed by 30 residues located in the vicinity of conserved motifs or the pocket (Fig. 4A). Eleven of these residue positions have been shown to significantly impact the basal activity of class A GPCR upon mutation (Fig. S3, Table S7). Although the conserved motifs themselves are important, they turned out to contribute little to the variation in basal activity, likely due to their low variance in the sequence. It may be expected that variations around these motifs have a strong impact on the basal activity. Interestingly, the rest of the 30 residues are in or around the odorant-binding pocket, indicating that ligand-dependent and -independent activities are intertwined. This is consistent with our previous finding that ORs with high basal activity respond to broad spectra of odorants and *vice versa*^21^. We found 5 residues important for both the basal activity and the ligand specificity/selectivity (Fig. 4), 4 of which (Ballesteros-Weinstein numbers 2.54, 3.36, 3.38 and 7.42) are in the so-called “connector” region of the GPCR TM bundle. These residues might mediate the interplay between the agonist-induced and the basal activations, in line with the role of the connector in GPCR activation^38^.

The machine learning approach captured complex information, going beyond a simple sequence similarity analysis to identify ligands (Fig. S5). The protocol is readily applicable to the entire mammalian OR families and will significantly accelerate the decoding of combinatorial OR codes for odors. It is an open loop where newly identified OR-odorant pairs and relevant residues can be added to continuously improve the next-generation model. Here, the ligand with the least data was associated with ~60 ORs. Note that good initial guesses are important, since it is unfeasible to test all the possible subsets of so many residues using a brute force approach. Owing to the large amount of data, i.e. *in vitro* functional assays, site-directed mutagenesis, GPCR structures and sequences, as well as molecular modeling, we could generate heuristics to decipher how nature has encoded specific functions of ORs into their varied sequences. The approach is also applicable to other protein families and functions, provided that enough data is available to generate heuristics and to train the machine learning models. The discovery of residue subsets associated with given functions may hint to evolutionary hotspots and compensate existing tools such as phylogenetic analysis based on full sequences.

## Materials and Methods

### Chemicals and OR constructs

Odorants were purchased from Sigma Aldrich. They were dissolved in DMSO to make stock solutions at 1 mM then diluted freshly in optimal MEM (ThermoFisher) to prepare the odorant stimuli. The OR constructs were kindly provided by Dr. Hanyi Zhuang (Shanghai Jiaotong University, China). Site-directed mutants were constructed using the Quikchange site-directed mutagenesis kit (Agilent Technologies). The sequences of all plasmid constructs were verified by both forward and reverse sequencing (Sangon Biotech, Shanghai, China).

### Cell culture and transfection

We used Hana3A cells, a HEK293T-derived cell line that stably expresses receptor-transporting proteins (RTP1L and RTP2), receptor expression-enhancing protein 1 (REEP1) and olfactory G protein (Gα_olf_)^39^ Th cells were grown in MEM (Corning) supplemented with 10% (vol/vol) fetal bovine serum (FBS) (ThermoFisher), added with 100 μg/ml penicillin-streptomycin (ThermoFisher), 1.25 μg/ml amphotericin (Sigma Aldrich), and 1 μg/ml puromycin (Sigma Aldrich).

All constructs were transfected into the cells using Lipofectamine 2000 (ThermoFisher). Before the transfection, the cells were plated on 96-well plates (NEST) and incubated overnight in MEM with 10% FBS at 37 °C and 5% CO_2_. For each 96-well plate, 2.4 μg of pRL-SV40, 2.4 μg of CRE-Luc, 2.4 μg of mouse RTP1S, and 12 μg of receptor plasmid DNA were transfected. The cells were subjected to Luciferase assay 24 hours after transfection.

### Luciferase assay

Luciferase assay was performed with the Dual-Glo Luciferase Assay Kit (Promega) following the protocol in ref.^39^. OR activation triggers the Gα_olf_-driven AC-cAMP-PKA signaling cascade and phosphorylates CREB. Activated CREB induces luciferase gene expression, which can be quantified luminometrically (measured here with a bioluminescence plate reader (MD SPECTRAMAX L)). Cells were co-transfected with *firefly* and *Renilla* luciferases where *firefly* luciferase served as the cAMP reporter. *Renilla* luciferase is driven by a constitutively active simian virus 40 (SV40) promoter (pRL-SV40; Promega), which served as a control for cell viability and transfection efficiency. The ratio between *firefly* luciferase versus *Renilla* luciferase was measured. Normalized OR activity is calculated as (*L*_N_ – *L*_min_)/(*L*_max_ –*L*_min_), where *L*_N_ is the luminescence in response to the odorant, and *L*_min_ and *L*_max_ are the minimum and maximum luminescence values on a plate, respectively. The assay was carried out as follows: 24h after transfection, medium was replaced with 100 μL of odorant solution (at different doses) diluted in Optimal MEM (ThermoFisher), and cells were further incubated for 4 h at 37 °C and 5% CO_2_. After incubation in lysis buffer for 15 minutes, 20 μl of Dual-Glo™ Luciferase Reagent was added to each well of 96-well plate and *firefly* luciferase luminescence was measured. Next, 20 μl Stop-Glo Luciferase Reagent was added to each well and *Renilla* luciferase luminescence was measured. Data analysis followed the published procedure in ref.^39^. Three-parameter dose-response curves were fit with GraphPad Prism.

### Molecular modeling

Homology modeling and MD simulations were carried out using previously established procedures (see Supplementary Materials and Methods).

### Machine learning

Data of OR response to the four odorants were collected from the literature. ORs that responded at high concentrations but not in a clear dose-dependent manner were considered ambiguous and discarded. OR sequence alignment was from our previous work^18^. Residue positions that contain gaps were removed. The sequences were digitized by assigning three physicochemical scores (features) per residue^40^. We removed features that showed low variance (< 10%) or high correlation (Pearson’s correlation coefficient > 0.85). Then, 3 steps of feature selection were applied (Fig. 1A): i) rational residue selection based on molecular modeling and site-directed mutagenesis; ii) Relief-F filter^41^ to identify features that are most relevant to partitioning the given problem; and iii) simulated annealing (SA)^42^, a global search that produces random perturbations to the selected features by substituting one of them with another from the unselected ones. At each SA iteration, the model performance was evaluated to decide whether to accept the perturbation. The SA was iterated until convergence to obtain an optimized selection of features.

RF, RPART and SVM classifiers were built using the Caret package in R^42^. Highly imbalanced data (i.e. the classes have very different sizes) were handled by ROSE^43^. The data were split into 80% training set and 20% test set. Parameters tuning was carried out by 5-fold cross validation repeated 5 times on the training set, followed by assessment on the validation set to choose the best models. The same procedure was employed to build the final models using all the data.

Matthew’s correlation coefficient (MCC)^27^ and Cohen’s Kappa^28^ were used as metrics of predictivity for the 2-class and the 3-class classifiers, respectively. Both metrics are generally considered more informative than other confusion matrix measures (e.g. accuracy, precision, recall and F1 score), especially for highly imbalanced data like the case here. MCC returns a value between −1 and +1, where +1 indicates a perfect prediction, 0 at a random prediction, and −1 at an inverse prediction. Cohen’s kappa returns a value <= 1, where 1 means perfect prediction and 0 or negative values indicate no agreement or useless models.

## General

We acknowledge GENCI-CINES (France) for providing access to the supercomputer OCCIGEN (grant A0040710477 to J.G). We also thank Dr. Sébastien Fiorucci, Dr. Jérémie Topin, Ms. Priyanka Meesa and Mr. Shiyi Jiang for critical reading and editing the manuscript.

## Author contributions

X.C, Y.Y and J.G designed research; X.C, W.R, J.P, C.A.D.M, and L.X performed research; X.C, W.R, J.P, C.A.D.M, L.X, H.M, Y.Y and J.G analyzed data; and X.C, Y.Y and J.G wrote the paper. All authors have given approval to the final version of the manuscript.

