## Supporting Information for "Functions of olfactory receptors are decoded from their sequence"

**Supplementary Materials**

### Supplementary Materials and Methods

*Homology modeling and docking*. Homology models of mOR256-3, mOR256-8 and mOR256-31 were built using the approach in ref. [1,2]. Four X-ray crystal structures of class A GPCRs were used as templates, Rhodopsin (1U19), CXCR4 (3ODU), A2aR (2YDV) and CXCR1 (2LNL), to build 100 models with Modeller v9.15 [3]. For docking, we chose the model with the lowest DOPE score. Autodock Vina [4] and the Haddock 2.2 webserver [5] were used to identify a common top-ranked binding pose for each odorant. Residues in the putative ligand-binding pocket were set flexible during docking.

*Molecular dynamics*. The receptor-odorant complexes were embedded in a bilayer of POPC using Desmond-Maestro (v2016.1, non-commercial distribution) [6]. Each system was solvated in a periodic 75 × 75 × 105 $Å^3$ box of explicit water and neutralized with 0.15 M of $Na^+$ and $Cl^-$ ions. Effective point charges of the ligands were obtained by RESP fitting [7] of the electrostatic potentials calculated with the HF/6-31G* basis set using Gaussian 09 [8]. The Amber 99SB-ildn [9], lipid 14 [10] and GAFF [11] force fields were used for the proteins, the lipids and the ligands, respectively. The TIP3P [12] and the Joung-Cheatham [13] models were used for the water and the ions, respectively.

After energy minimization, all-atom MD simulations were carried out using Gromacs 5.1 patched with the PLUMED 2.3 plugin [14]. Each system was gradually heated to 310 K and pre-equilibrated during 10 ns of brute-force MD in the *NPT*-ensemble (see SI-Methods for details). The replica exchange with solute scaling (REST2) [15] technique was then employed to enhance the sampling with 48 replicas in the *NVT* ensemble. The protein and the ligands were considered as “solute” in the REST2 scheme–force constants of their van der Waals, electrostatic and dihedral terms were subject to scaling. The effective temperatures used for generating the REST2 scaling factors ranged from 310 K to 700 K, following a distribution calculated with the Patriksson-van der Spoel approach [16]. Exchange between replicas was attempted every 1000 simulation steps. This setup resulted in an average exchange probability of ~40%. The original unscaled replica (at 310 K effective temperature) was collected and analyzed. The first 10 ns were discarded for equilibration. Cluster analysis of the ligand binding pose was carried out on the non-restrained trajectory using g_cluster in Gromacs tools with the GROMOS method [17]. The middle structure of the most populated cluster was selected as the final binding pose.

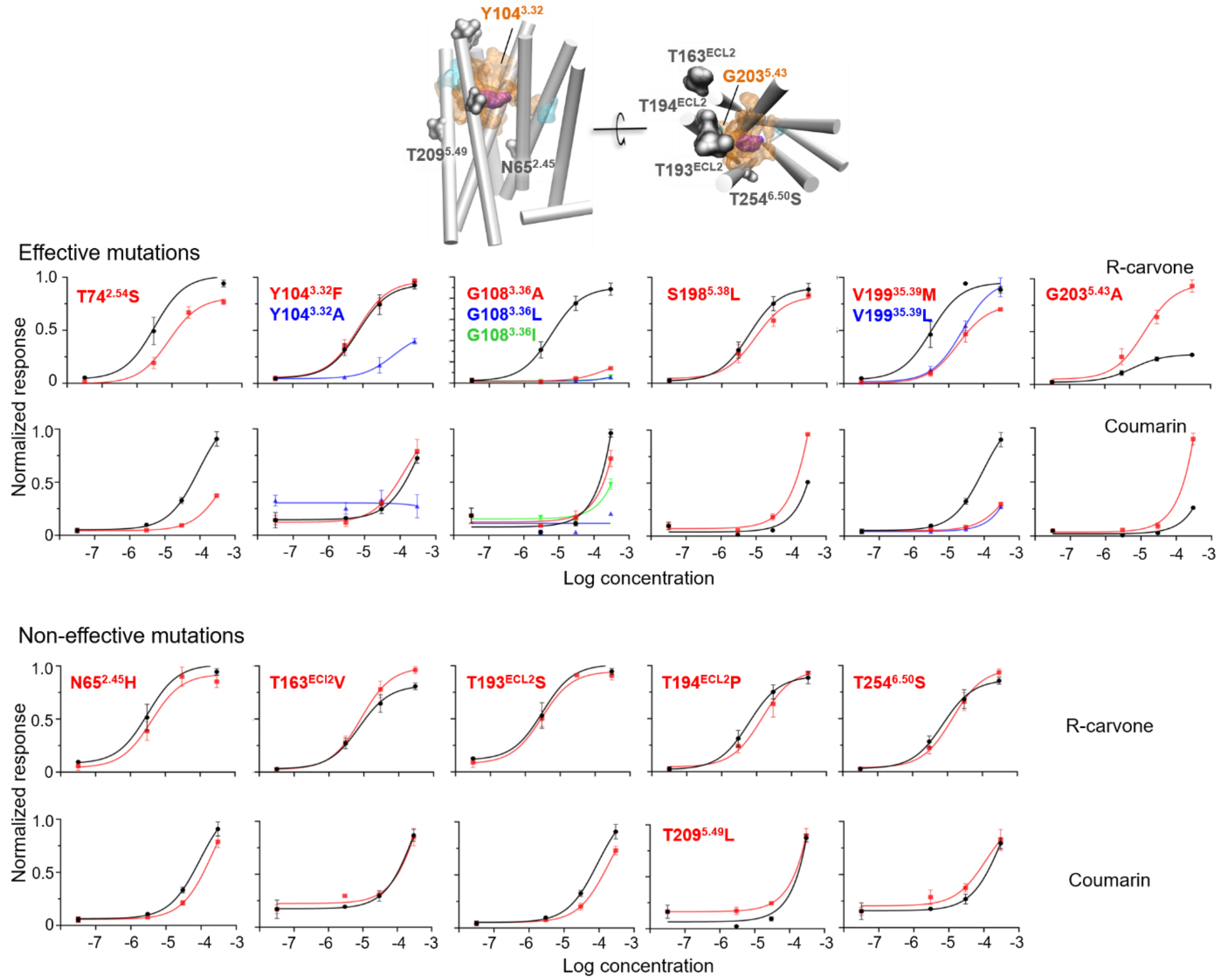


**Fig. S1. Dose-dependent response of mOR256-31 mutants to coumarin and R-carvone.** The residues were mutated to the corresponding ones in mOR256-8 (red), a narrowly tuned OR that does not response to R-carvone or coumarin [18]. Four other mutations (blue and green) were made to Y104$^{3.32}$, G108$^{3.36}$ and V199$^{5.39}$.

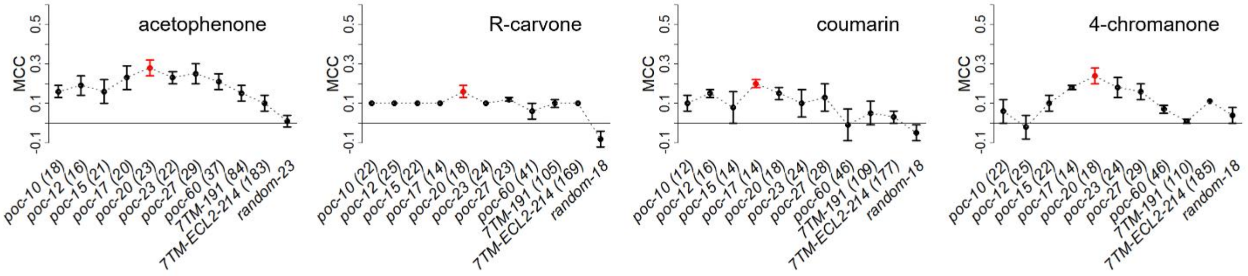


**Fig. S2.** Matthew's correlation coefficient (MCC) as metric of RF classifier predictivity, tested on the external test set of 43 ORs. Labeled in parenthesis is the number of optimal residues that gave the best performance in each model. The best models are highlighted in red. The control models (random) were built with randomly chosen residues (same number of residues as in the best model).

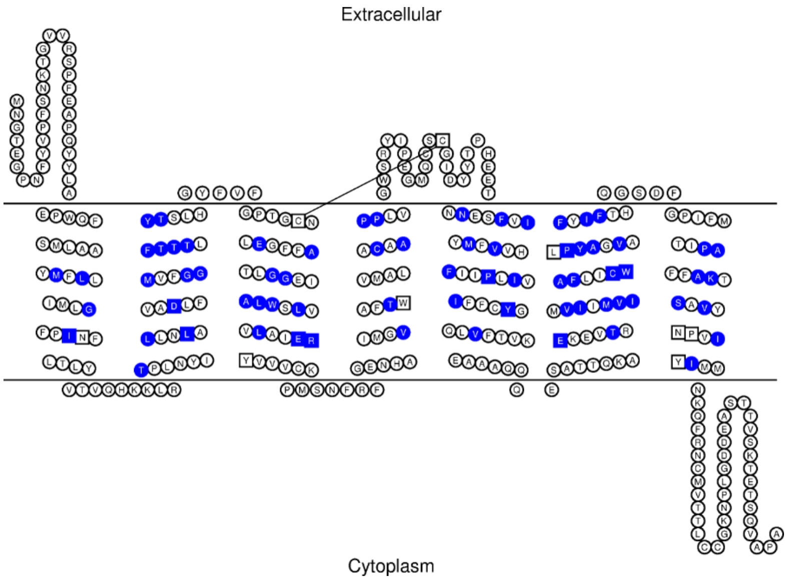


**Figure S3.** Residue positions that, upon mutation, significantly altered the basal activity of class A GPCRs (Table S8).

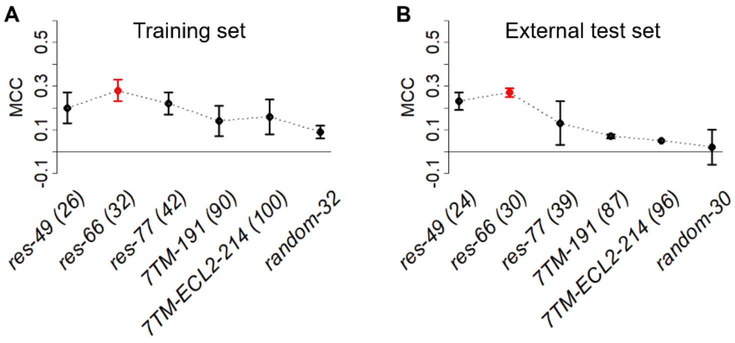


**Fig. S4. RPART classifiers' performance on OR basal activity.** (A) The training set during 5-fold cross validation repeated 5 times. (B) The external test set repeated 5 times. Labeled in parenthesis is the number of optimal residues that gave the best performance in each model. The best models are highlighted in red. The control models (random) were built with randomly chosen residues (same number of residues as in the best model).

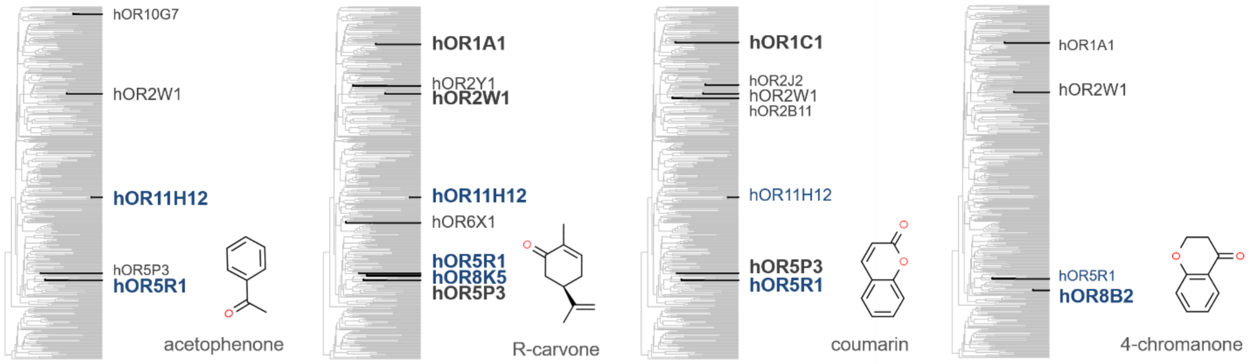


**Figure S5.** Current knowledge of the combinatorial codes for the 4 odorants illustrated on the phylogenetic tree of hORs. The large to small hOR label sizes indicate the strength of response (EC50 < 10 μM, 10 μM < EC50 < 100 μM and EC50 > 100 μM, respectively). hORs identified in this work are labelled in blue.

**Table S1. Deorphanized human and mouse ORs to date.**

| | | Count | Refs |
|---|---|---|---|
| Human ORs | hOR10A5, hOR10A6, hOR10A7, hOR10AG1, hOR10C1, hOR10G3, hOR10G4, hOR10G6, hOR10G7, hOR10H1, hOR10J3, hOR10J5, hOR10K1, hOR11A1, hOR11G2, hOR11H1, hOR11H13, hOR11H2, hOR11H4, hOR11H6, hOR11H7, hOR13C2, hOR13C5, hOR13C8, hOR13C9, hOR13G1, hOR14C36, hOR14J1, hOR14L1, hOR1A1, hOR1A2, hOR1C1, hOR1D2, hOR1D5, hOR1E1, hOR1E3, hOR1G1, hOR1J1, hOR1J4, hOR1L3, hOR1N1, hOR1N2, hOR1S1, hOR1S2, hOR2A25, hOR2A4, hOR2A7, hOR2AG1, hOR2AK2, hOR2AT4, hOR2B11, hOR2B2, hOR2B3, hOR2C1, hOR2C3, hOR2D2, hOR2F1, hOR2G2, hOR2J2, hOR2J3, hOR2L2, hOR2L5, hOR2L8, hOR2M3, hOR2M4, hOR2M7, hOR2T1, hOR2T10, hOR2T11, hOR2T29, hOR2T34, hOR2T35, hOR2T6, hOR2T8, hOR2V2, hOR2W1, hOR2Y1, hOR3A1, hOR3A3, hOR3A4, hOR4A16, hOR4A4, hOR4A47, hOR4A5, hOR4D1, hOR4D10, hOR4D2, hOR4D6, hOR4D9, hOR4E2, hOR4F17, hOR4F4, hOR4K1, hOR4K14, hOR4K15, hOR4K5, hOR4M1, hOR4N2, hOR4N5, hOR4P4, hOR4Q3, hOR4X2, hOR51A2, hOR51B2, hOR51B4, hOR51B5, hOR51B6, hOR51D1, hOR51E1, hOR51E2, hOR51F1, hOR51I2, hOR51J1, hOR51L1, hOR51M1, hOR51S1, hOR51T1, hOR51V1, hOR52A5, hOR52D1, hOR52E5, hOR52E8, hOR52H1, hOR52I1, hOR52J3, hOR52K1, hOR52K2, hOR52L1, hOR52M1, hOR52N1, hOR52N2, hOR52N5, hOR52R1, hOR52W1, hOR56A3, hOR56A4, hOR56A5, hOR56B4, hOR5A1, hOR5A2, hOR5AK2, hOR5AL1, hOR5AN1, hOR5AS1, hOR5B12, hOR5B17, hOR5B3, hOR5C2, hOR5D14, hOR5D18, hOR5H2, hOR5J2, hOR5K1, hOR5L1, hOR5M11, hOR5P2, hOR5P3, hOR6C1, hOR6C65, hOR6C68, hOR6J1, hOR6K2, hOR6P1, hOR6S1, hOR7A5, hOR7C1, hOR7D4, hOR7G2, hOR8B12, hOR8B2, hOR8B3, hOR8D1, hOR8G5, hOR8J1, hOR8K3, hOR9A2, hOR9G1, hOR9G4, hOR9Q1, hOR9Q2 | 180 | [19-34] |
| Mouse ORs | Olfr64, Olfr1019, Olfr1032, Olfr1062, Olfr1062, Olfr1079, Olfr1079, Olfr109, Olfr109, Olfr1093, Olfr1104, Olfr1104, Olfr124, Olfr1264, Olfr1324, Olfr1325, Olfr1328, Olfr1341, Olfr1352, Olfr1356, Olfr1364, Olfr1370, Olfr1377, Olfr1377, Olfr1395, Olfr1411, Olfr1413, Olfr145, Olfr1484, Olfr15, Olfr1509, Olfr151, Olfr1512, Olfr154, Olfr16, Olfr16, Olfr160, Olfr161, Olfr167, Olfr167, Olfr168, Olfr171, Olfr175, Olfr19, Olfr2, Olfr202, Olfr211, Olfr214, Olfr221, Olfr24, Olfr311, Olfr320, Olfr323, Olfr340, Olfr406, Olfr429, Olfr447, Olfr459, Olfr476, Olfr480, Olfr49, Olfr491, Olfr50, Olfr502, Olfr508, Olfr510, Olfr514, Olfr532, Olfr544, Olfr545, Olfr547, Olfr549, Olfr554, Olfr556, Olfr556, Olfr558, Olfr56, Olfr569, Olfr599, Olfr609, Olfr61, Olfr611, Olfr616, Olfr62, Olfr620, Olfr638, Olfr642, Olfr644, Olfr65, Olfr653, Olfr661, Olfr67, Olfr672, Olfr677, Olfr678, Olfr68, Olfr683, Olfr685, Olfr690, Olfr691, Olfr715, Olfr73, Olfr74, Olfr744, Olfr749, Olfr749, Olfr790, Olfr796, Olfr876, Olfr876, Olfr889, Olfr895, Olfr90, Olfr909, Olfr919, Olfr961, Olfr963, Olfr978, Olfr979, Olfr979, Olfr982, Olfr983, Olfr992, Olfr992 | 124 | [19,21,22,33,35-60] |

**Table S2. OR-odorant pairs used in machine learning model training and cross validation.**

| | Responsive | Non-responsive | Ref |
|---|---|---|---|
| acetophenone | hOR10G7, hOR2W1, hOR5P3, Olfr1009, Olfr1044, Olfr1047, Olfr1054, Olfr1062, Olfr1079, Olfr109, Olfr1093, Olfr1094, Olfr110, Olfr1104, Olfr1126, Olfr1170, Olfr1238, Olfr124, Olfr133, Olfr1333, Olfr1352, Olfr136, Olfr1370, Olfr1377, Olfr143, Olfr1443, Olfr1444, Olfr1448, Olfr145, Olfr1463, Olfr1469, Olfr1484, Olfr151, Olfr156, Olfr160, Olfr166, Olfr167, Olfr168, Olfr191, Olfr196, Olfr201, Olfr203, Olfr205, Olfr30, Olfr339, Olfr346, Olfr347, Olfr376, Olfr414, Olfr429, Olfr430, Olfr434, Olfr466, Olfr476, Olfr478, Olfr490, Olfr491, Olfr494, Olfr502, Olfr51, Olfr556, Olfr57, Olfr60, Olfr62, Olfr736, Olfr739, Olfr744, Olfr746, Olfr749, Olfr874, Olfr875, Olfr876, Olfr887, Olfr888, Olfr889, Olfr890, Olfr895, Olfr904, Olfr920, Olfr923, Olfr935, Olfr983 | hOR10A3, hOR10A7, hOR10C1, hOR10J5, hOR10K1, hOR10Q1, hOR10R2, hOR10X1, hOR12D2, hOR13D1, hOR14J1, hOR14L1, hOR1C1, hOR1D5, hOR1E2, hOR1G1, hOR1J2, hOR1L8, hOR1N2, hOR2A12, hOR2A4, hOR2AG1, hOR2AT4, hOR2G6, hOR2L13, hOR2L2, hOR2L3, hOR2L8, hOR2M4, hOR2M7, hOR2T5, hOR2W5, hOR4A16, hOR4C15, hOR4C3, hOR4C5, hOR4F16, hOR4M2, hOR4N4, hOR4S1, hOR51E1, hOR51E2, hOR51F1, hOR51L1, hOR52B2, hOR52D1, hOR52I1, hOR56A1, hOR5AL1, hOR5D16, hOR5H1, hOR5H2, hOR5H6, hOR5M9, hOR5P2, hOR5T1, hOR6C3, hOR6C6, hOR6C65, hOR6C75, hOR6F1, hOR6K3, hOR6K6, hOR6N2, hOR7A10, hOR7A5, hOR7G2, hOR8A1, hOR8B8, hOR8G5, hOR9A2, hOR9Q2, Olfr1019, Olfr1264, Olfr1324, Olfr1395, Olfr15, Olfr171, Olfr174, Olfr19, Olfr202, Olfr221, Olfr311, Olfr323, Olfr340, Olfr508, Olfr532, Olfr554, Olfr558, Olfr569, Olfr599, Olfr609, Olfr61, Olfr611, Olfr632, Olfr638, Olfr64, Olfr653, Olfr67, Olfr683, Olfr685, Olfr715, Olfr796, Olfr979, Olfr992 | 21,61-63 |
| R-carvone | hOR1A1, hOR2W1, hOR2Y1, hOR5P3, hOR6X1, Olfr1079, Olfr124, Olfr1352, Olfr1356, Olfr168, Olfr174, Olfr263, Olfr340, Olfr362, Olfr45, Olfr50, Olfr556, Olfr895, Olfr992 | hOR10A2, hOR10A5, hOR10A6, hOR10A7, hOR10AG1, hOR10C1, hOR10G6, hOR10G7, hOR10K2, hOR10Q1, hOR10X1, hOR11H1, hOR12D3, hOR13C2, hOR13C4, hOR13C8, hOR13D1, hOR14A2, hOR1C1, hOR1D5, hOR1E2, hOR1G1, hOR1J1, hOR1L8, hOR1N1, hOR1N2, hOR1S1, hOR1S2, hOR2A1, hOR2A12, hOR2AG1, hOR2AT4, hOR2B2, hOR2C1, hOR2D2, hOR2F1, hOR2G6, hOR2J2, hOR2K2, hOR2L2, hOR2L5, hOR2L8, hOR2M3, hOR2M7, hOR2T29, hOR2T5, hOR2T8, hOR2Z1, hOR4A16, hOR4A5, hOR4B1, hOR4C11, hOR4C15, hOR4C3, hOR4D10, hOR4D5, hOR4F16, hOR4F17, hOR4F4, hOR4F5, hOR4K13, hOR4K2, hOR4K3, hOR4K5, hOR4M1, hOR4N2, hOR4N4, hOR4N5, hOR4S1, hOR4X2, hOR5A2, hOR5AK3, hOR5AN1, hOR5AR1, hOR5B12, hOR5B17, hOR5D16, hOR5H6, hOR5L1, hOR5M11, hOR5M9, hOR5T2, hOR5V1, hOR6A2, hOR6C65, hOR6C75, hOR6F1, hOR6K6, hOR6M1, hOR6P1, hOR7A5, hOR7C1, hOR7D4, hOR7G1, hOR7G2, hOR7G3, hOR8A1, hOR8B12, hOR8K3, hOR8G5, hOR8U8, hOR9G1, hOR9Q2, Olfr1019, Olfr1062, Olfr109, Olfr1104, Olfr1264, Olfr1324, Olfr1341, Olfr1377, Olfr1395, Olfr15, Olfr171, Olfr19, Olfr202, Olfr221, Olfr311, Olfr323, Olfr429, Olfr532, Olfr554, Olfr558, Olfr569, Olfr599, Olfr609, Olfr61, Olfr611, Olfr632, Olfr638, Olfr64, Olfr653, Olfr67, Olfr683, Olfr685, Olfr715, Olfr749, Olfr796, Olfr979, Olfr983 | 20,21,35,63,64 |
| coumarin | hOR1C1, hOR2B11, hOR2J2, hOR2W1, hOR5P3, Olfr1062, Olfr109, Olfr1104, Olfr124, Olfr1328, Olfr1341, Olfr1352, Olfr1377, Olfr263, Olfr508, Olfr556, Olfr749, Olfr876, Olfr895, Olfr983 | hOR1A1, hOR10J5, hOR2C1, hOR2M7, Olfr167, Olfr168, Olfr992, Olfr15, Olfr171, Olfr340, Olfr429, Olfr715, Olfr1395, Olfr1324, Olfr323, Olfr1264, Olfr979, Olfr796, Olfr174, Olfr202, Olfr61, Olfr532, Olfr311, Olfr1019, Olfr221, Olfr19, Olfr447, Olfr514, Olfr64, Olfr632, Olfr569, Olfr685, Olfr609, Olfr611, Olfr683, Olfr653, Olfr638, Olfr554, Olfr67, Olfr599, Olfr558 | 18,19,21,63 |
| 4-chromanone | hOR1A1, hOR2W1, Olfr1062, Olfr1079, Olfr1104, Olfr1352, Olfr1377, Olfr1395, Olfr167, Olfr168, Olfr171, Olfr311, Olfr340, Olfr556, Olfr895, Olfr983 | hOR5P3, hOR2J2, hOR10J5, hOR2C1, hOR2M7, Olfr749, Olfr109, Olfr508, Olfr1341, Olfr992, Olfr15, Olfr429, Olfr715, Olfr1324, Olfr323, Olfr1264, Olfr979, Olfr796, Olfr174, Olfr202, Olfr61, Olfr532, Olfr1019, Olfr221, Olfr19, Olfr447, Olfr514, Olfr64, Olfr632, Olfr569, Olfr685, Olfr609, Olfr611, Olfr683, Olfr653, Olfr638, Olfr554, Olfr67, Olfr599, Olfr558 | 21 |

**Table S3. mOR256-31 residues in direct contact with the odorants.**

| Ballesteros-Weinstein nomenclature | Occupation (%) during MD [a] | | | Literature data (mutagenesis) | |
|---|---|---|---|---|---|
| | Coumarin | R-carvone | Acetophenone | Receptor [b] | Refs. |
| 2.53 | 100 | 38 | 44 | | |
| 3.29 | 0 | 72 | 49 | hOR1A1 | |
| 3.30 | 0 | 17 | 0 | mOR256-3, mOR256-8 | |
| 3.32 | 100 | 100 | 96 | hOR1A1, hOR2AG1, mOR-EG, mOR256-3 | |
| 3.33 | 100 | 100 | 100 | hOR1A1, mOR-EG, mOR42-3 | |
| 3.36 | 100 | 99 | 99 | hOR1A1, hOR1A2, mOR-EG, mOR42-3, mOR256-3 | |
| 3.37 | 9 | 81 | 97 | hOR1A1, hOR1A2, mOR42-3, mOR244-3 | |
| 3.38 | 0 | 11 | 1 | | [1,2,18,35 |
| 5.42 | 40 | 92 | 99 | hOR1A1, mOR-EG, mOR42-3 | ,39,65-69] |
| 5.43 | 0 | 74 | 99 | mOR42-3, mOR256-3 | |
| 5.46 | 94 | 71 | 59 | hOR1A1, hOR1A2, hOR2AG1, mOR-EG, mOR42-3 | |
| 5.47 | 0 | 89 | 97 | hOR1A1, mOR-EG | |
| 6.48 | 100 | 85 | 97 | hOR1A1, hOR7D4, mOR256-3 | |
| 6.51 | 100 | 100 | 73 | hOR1A1, hOR1A2, mOR-EG, mOR42-3 | |
| 6.52 | 0 | 96 | 100 | | |
| 6.55 | 0 | 97 | 97 | mOR-EG | |
| 7.42 | 100 | 30 | 1 | hOR1A1, hOR1A2, hOR2AG1, mOR-EG | |

[a] Percentage of the simulation trajectory where the residue was within 5 Å distance of the odorants.

[b] Receptor response to odorants was affected upon mutation.

**Table S4.** Six subsets of residues tested in machine learning of OR response to odorants.

| Subset | Number of residues | Ballesteros-Weinstein nomenclature |
|---|---|---|
| *poc-17* | 17 | 2.53 3.29 3.30 3.32 3.33 3.36 3.37 3.38 5.42 5.43 5.46 5.47 6.48 6.51 6.52 6.55 7.42 |
| *poc-20* | 20 | 2.53 2.54 3.29 3.30 3.32 3.33 3.36 3.37 3.38 5.38 5.39 5.42 5.43 5.46 5.47 6.48 6.51 6.52 6.55 7.42 |
| *poc-27* | 27 | 2.53 2.54 2.57 3.29 3.30 3.31 3.32 3.33 3.35 3.36 3.37 3.38 4.48 4.52 5.38 5.39 5.41 5.42 5.43 5.46 5.47 6.48 6.51 6.52 6.55 7.39 7.42 |
| *poc-60* | 60 | 2.53 2.54 2.57 2.61 3.25 3.26 3.27 3.28 3.29 3.30 3.31 3.32 3.33 3.34 3.35 3.36 3.37 3.38 3.39 3.40 3.41 4.52 4.53 4.56 4.57 4.60 5.35 5.36 5.37 5.38 5.39 5.40 5.41 5.42 5.43 5.44 5.45 5.46 5.47 5.48 5.49 5.50 5.51 6.44 6.45 6.47 6.48 6.49 6.50 6.51 6.52 6.53 6.54 6.55 6.56 7.35 7.38 7.39 7.41 7.42 |
| *7TM-191* | 191 | 1.32–1.65, 2.39–2.67, 3.25–3.55, 4.40–4.61, 5.37–5.65, 6.32–6.56, 7.35–7.55 |
| *7TM-ECL2-214* | 214 | Subset V + 23 residues between the 2 conserved cysteines in ECl2 |

**Table S5**. Newly identified OR-odorant pairs and EC50 (with 95% confidence interval) [a] in Hana3A cells.

| | acetophenone | R-carvone | coumarin | 4-chromanone |
|---|---|---|---|---|
| hOR11H12 | 4.7 (0.7–29.9) | 16.3 (5.5–48.4) | 91.9 (16.6–506.7) | n.r. |
| hOR5R1 | 3.4 (0.9–12.7) | 12 (3.2–32.7) | 10.2 (2.2–64.3) | 217.8 (10.5–4506) |
| hOR8B2 | n.r. | n.r. | n.r. | 15 (2.2–101.4) |
| hOR8K5 | n.r. | 4.5 (1.0–21.3) | n.r. | n.r. |
| Olfr1097 | 57.3 (14.5–226.8) | n.r. | 7.2 (1.0–54.2) | 9.5 (1.8–49.5) |
| Olfr1016 | n.r. | n.r. | 72.4 (14.3–336.1) | n.r. |
| Olfr1057 | n.r. | n.r. | 3.8 (1.3–11.5) | 5.8 (2.6–13.2) |
| Olfr1156 | 46.2 (9.6–222.8) | 30.5 (1.2–789) | n.r. | n.r. |
| Olfr285 | n.r. | n.r. | 66.7 (32.3–137.8) | 72.8 (14.5–366.7) |
| Olfr924 | n.r. | n.r. | 28 (4.5–173) | n.r. |

[a] n.r. indicates no significant response up to 1 mM.

**Table S6.** Residue positions that, upon mutation, significantly altered the basal activity of class A GPCRs.

| Receptor | Gain of function | Loss of function | Ref. |
|---|---|---|---|
| A1b adrenergic receptor | 3.49, 3.50, 6.30, 6.34 | | 70,71 |
| Adenosine 1a receptor | 6.34 | | 71 |
| Adenosine 2a receptor | 3.39 | 4.46, 4.49, 4.56, 4.58, 4.60, 4.61, 5.35, 5.38, 5.42, 5.40, 5.47 | 72,73 |
| Angiotensin II type 1 receptor | | 3.35, 3.41, 4.49 | 74,75 |
| B2 adrenergic receptor | 3.49, 6.30, 6.44, 6.47, 7.46 | | 76-78 |
| B3 adrenergic receptor | | 5.47, 6.51, 6.52 | 79 |
| Canabinoid receptor 1 | 2.37, 2.63, 3.43 | | 80,81 |
| C-C chemokine receptor 5 | 2.65 | | 82 |
| Cholecystokinin B receptor | 3.32, 5.40, 5.52 | | 83 |
| C-X-C chemokin receptors 1–4 | 2.53, 2.56, 3.35, 6.40 | | 84-86 |
| δ-opioid receptor | 3.32, 7.42, 8.53 | | 87,88 |
| Follicle-stimulating hormone receptor | 3.32, 5.54, 6.30 | | 89 |
| GPR18 | | 3.39 | 90 |
| Growth hormone secretagogue receptor | 3.40, 3.43 | 6.48 | 91 |
| Gonadotropin-releasing hormone receptor | 2.53, 6.40, 6.47, 6.52 | | 92,93 |
| Histamine 1 receptor | 6.40 | | 94 |
| Luteinizing hormone receptor | 1.41, 1.46, 2.43, 3.43, 5.54, 6.30, 6.34, 6.37, 6.38, 6.41, 6.43, 6.44, 6.47, 6.48 | 2.50, 5.58 | 89,95 |
| µ-opioid receptor | 3.35, 3.49 | | 96,97 |
| Muscarinic receptors 1–5 | 6.58, 6.59 | | 98 |
| Neurotensin receptor 1 | 7.42 | | 99 |
| Nociceptin receptor | 3.35 | | 100 |
| Parathyroid hormone 1 receptor | 2.43, 6.37, 7.52 | | 101 |
| P2Y12 receptor | | 3.50 | 102 |
| Rhodopsin | 2.46, 2.57, 2.61, 3.28, 6.40, 7.38, 7.39, 7.42, 7.43, 7.54 | | 89,103-107 |
| Serotonin 5HT4 receptor | | 2.50, 3.36, 6.48 | 108,109 |
| Serotonin 5HT2a receptor | 6.34 | | 110 |
| Thyrotropin receptor | 1.39, 1.49, 2.43, 2.53, 2.56, 2.58, 3.32, 3.36, 3.43, 3.40, 5.44, 5.54, 5.58, 6.30, 6.34, 6.37, 6.40, 6.42, 6.43, 6.44, 6.48, 6.50, 6.52, 6.54, 6.56, 6.59, 7.42, 7.45, 7.47, 7.52 | 2.59, 2.60, 5.50, 6.47 | 89,101,111-114 |
| Vasopressin receptor 2 | 3.43, 3.50, 5.62 | | 101 |
